# Efficient breeding of industrial brewing yeast strains using CRISPR/Cas9-aided mating-type switching

**DOI:** 10.1101/2021.07.05.450511

**Authors:** Kristoffer Krogerus, Eugene Fletcher, Nils Rettberg, Brian Gibson, Richard Preiss

**Author notes:** Address correspondence to Kristoffer Krogerus.

## Abstract

Yeast breeding is a powerful tool for developing and improving brewing yeast in a number of industry-relevant respects. However, breeding of industrial brewing yeast can be challenging, as strains are typically sterile and have large complex genomes. To facilitate breeding, we used the CRISPR/Cas9 system to generate double-stranded breaks in the *MAT* locus, generating transformants with a single specified mating type. The single mating type remained stable even after loss of the Cas9 plasmid, despite the strains being homothallic, and these strains could be readily mated with other brewing yeast transformants of opposite mating type. As a proof of concept, we applied this technology to generate yeast hybrids with an aim to increase β-lyase activity for fermentation of beer with enhanced hop flavour. First, a genetic and phenotypic pre-screening of 38 strains was carried out in order to identify potential parent strains with high β-lyase activity. Mating-competent transformants of eight parent strains were generated, and these were used to generate over 60 hybrids that were screened for β-lyase activity. Selected phenolic off-flavour positive (POF+) hybrids were further sporulated to generate meiotic segregants with high β-lyase activity, efficient wort fermentation and lack of POF; all traits that are desirable in strains for the fermentation of modern hop-forward beers. Our study demonstrates the power of combining the CRISPR/Cas9 system with classic yeast breeding to facilitate development and diversification of brewing yeast.

**Key Points:**

- CRISPR/Cas9-based mating type switching was applied to industrial yeast strains
- Transformed strains could be readily mated to form intraspecific hybrids
- Hybrids exhibited heterosis for a number of brewing-relevant traits

## Introduction

The number of breweries and beer brands globally has expanded dramatically in recent decades (Garavaglia and Swinnen 2018). Consumers are also demanding higher product quality and beer with novel and diverse flavours (Aquilani et al. 2015; Carbone and Quici 2020; Gonzalez Viejo and Fuentes 2020). As much of beer flavour is yeast-derived (Holt et al. 2019), brewers may meet this demand and keep ahead of competition by diversifying their products through the use of different yeast strains. While a large and diverse range of yeast strains are naturally available, recent studies have shown that the vast majority of industrially used brewing strains group into one of two domesticated clades (Gallone et al. 2016; Gonçalves et al. 2016; Peter et al. 2018). These strains have evolved to efficiently ferment the complex sugars available in brewer’s wort. Non-brewing strains may therefore have difficulties completing fermentation in wort. Yeast breeding and hybridization has been shown to be a promising tool for developing and improving brewing yeast in a number of industry-relevant respects (Steensels et al. 2014; Krogerus et al. 2015; Mertens et al. 2015; Krogerus et al. 2016). It allows for the combination and enhancement of phenotypic traits from diverse sets of strains. Hybrids between *Saccharomyces cerevisiae* brewing strains and wild *Saccharomyces* strains, for example, have shown both efficient wort fermentation and a more diverse aroma profile (Mertens et al. 2015; Nikulin et al. 2018). However, brewing strains of *S. cerevisiae*, especially those in the ‘Beer 1’/’Ale beer’ group, are typically sterile, which impedes their use in yeast breeding (Gallone et al. 2016; De Chiara et al. 2020; Shimoi et al. 2020).

Yeast breeding relies on the formation and interaction of mating-competent cells (Herskowitz 1988; Neiman 2011; Merlini et al. 2013). Mating-competent cells may form when a diploid *MAT***a**/*MATα* cell undergoes meiosis and produces haploid spores of either *MAT***a** or *MATα* mating type. When two cells of opposite mating type come in contact with each other, they can undergo mating and cell fusion. Haploid cells may also switch mating type following the repair of a double-stranded break (DSB) created by the HO endonuclease at the mating type locus (Haber 2012). As the *HO* gene is repressed in diploid (or polyploid) cells heterozygous for the mating type locus (*MAT***a**/*MATα*), their mating type remains stable. Such cells can, however, on rare occasions undergo loss of heterozygosity at the mating type locus, which results in the formation of non-haploid mating-competent cells with a single mating type (Gunge and Nakatomi 1972; Hiraoka et al. 2000). This is often exploited for breeding of sterile strains, such as brewing strains, in a process called ‘rare mating’ (Krogerus et al. 2017). However, spontaneous loss of heterozygosity events occur at low frequencies (< 10^−4^) and parent strains require selection markers to allow selection of successful crosses. Obtaining hybrids with industrial brewing strains can therefore be challenging and time-consuming.

To overcome these limitations, a number of engineering techniques have been developed to facilitate breeding of sterile yeast strains. Alexander et al. (Alexander et al. 2016) describe a technique that can be used to force mating-type change in *MAT***a***/MATα* cells by transformation with a plasmid carrying the *HO* gene under the control of an inducible promoter and a drug-resistance marker. Fukuda et al. (Fukuda et al. 2016) describe another approach, where *MAT***a**/*MATα* cells are transformed with a plasmid carrying either the *a1* or *α2* gene from the mating-type locus together with drug-resistance markers with promoters specific to either the *MAT***a***1* or *MATα2* gene products. Recently, the CRISPR/Cas9 system was also used to force mating-type changes in diploid cells by creating DSBs in the mating-type locus using a Cas9 enzyme (Xie et al. 2018). As the approach was, to our knowledge, only tested on heterothallic (*ho)* laboratory strains with a maximum ploidy of two, we wished to explore whether it could be applied to industrial brewing strains, which are homothallic and aneuploid (often with DNA contents close to tetraploid).

In this study, we therefore applied the CRISPR-based mating-type switching process developed by Xie et al. (Xie et al. 2018) to industrial brewing strains in the hope of isolating variants with a stable single mating-type. Furthermore, we ultimately wanted to use these stable single mating-type variants to readily generate hybrids between industrial brewing strains. As a proof of concept, we aimed to generate yeast hybrids with increased β-lyase activity for fermentation of beer with enhanced hop flavour from released thiols. Recent studies have highlighted the important contribution of volatile thiols to fruity hop aroma in beer (Gros et al. 2012; Cibaka et al. 2017; Dennenlöhr et al. 2020). These compounds are present in minute amounts, but are still perceivable thanks to low odour thresholds (Holt et al. 2019). In addition to these free thiols, a large fraction of the total thiols in hops are found in glutathionylated or cysteinylated form (Gros et al. 2012; Roland et al. 2016). These conjugated thiols do not impact aroma by themselves, but may transfer to wort during the brewing process. Therein, a volatile thiol may be enzymatically released from the conjugated thiol through β-lyase activity (Roncoroni et al. 2011). Hence, a beer fermented with a yeast strain high in β-lyase activity is expected to contain higher levels of volatile thiols than one from a strain with low activity.

Here, a set of 38 *S. cerevisiae* strains were first phenotypically and genetically screened in order to identify potential parent strains with high β-lyase activity. We then generated mating-competent transformants of eight parent strains with either *MAT***a** or *MATα* mating-types. These were then used to generate over 60 hybrids that were screened for β-lyase activity. As multiple parent strains were POF+ (phenolic off-flavour positive), a selection of hybrids were further sporulated to generate meiotic segregants lacking the POF trait. A range of hybrid segregants were obtained with high β-lyase activity, efficient wort fermentation and lack of POF; all traits that are desirable in strains for the fermentation of modern hop-forward beers. Our study demonstrates the power of combining the CRISPR/Cas9 system with classic yeast breeding to facilitate development and diversification of brewing yeast.

## Materials and methods

### Yeast strains

A list of strains used in this study is available in Supplementary Table S1.

### High-throughput phenotypic assays

β-lyase activity was estimated by measuring growth on various nitrogen sources containing carbon-sulfur bonds: cysteine, s-methylcysteine and cys-4MMP (synthesized according to Howell et al. (2004)). Media contained 0.17% Yeast Nitrogen Base without (NH_4_)_2_SO_4_ and amino acids, 1% glucose, 0.01% pyridoxal 5-phosphate, and 15 mM of the above listed nitrogen sources. Growth assays were carried out in 96-well plates, with 145 *µ*L media per well. Wells were inoculated (to a starting OD600 value of 0.1) with 5 *µ*L of washed pre-culture suspended in water to an OD600 value of 3. Plates were sealed with a Breathe-Easy membrane (Sigma-Aldrich, Espoo, Finland), and incubated at 25 °C for one week. OD600 values were measured on a VarioSkan plate reader (Thermo Scientific, USA), while cysteine content of the growth media was estimated using DTNB (Ellman 1958).

The ability to produce phenolic off-flavour was estimated using the absorbance-based method described by Mertens et al. (2017).

Micro-scale wort fermentations were carried out in Greiner deep-well plates containing 700 *µ*L of 15 °P wort. Yeast was inoculated to a starting OD600 of 0.1 from washed pre-culture suspended in water to an OD600 value of 3. Fermentations were carried out for 4 days at 25 °C, after which the plates were centrifuged and the supernatant was analysed by HPLC for fermentable sugars and ethanol.

### Cas9 plasmid construction and yeast transformations

Plasmid construction was carried out using the plasmid pCC-036 as backbone (Rantasalo et al. 2018). pCC-036 contains yeast codon-optimized Cas9 expressed under *TDH3p*, guiding RNA (gRNA) expressed under *SNR52p*, and *hygR* for selection on hygromycin. The two gRNA protospacer sequences, GTTCTAAAAATGCCCGTGCT and CAAATCATACAGAAACACAG, were obtained from Xie et al. (2018), and target *MAT***a** and *MAT*α, respectively. A synthetic DNA fragment with the gRNA sequence was ordered from Integrated DNA Technologies (Leuven, Belgium) as a gBlock and introduced into the plasmid with restriction enzyme-based techniques (Thermo Scientific, Vantaa, Finland). The ligated plasmid was transformed into *E. coli* TOP10 by electroporation, and plasmid correctness was confirmed by Sanger sequencing.

Yeast transformations were performed using an optimized stationary phase transformation protocol (Tripp et al. 2013). Overnight yeast cultures were pelleted and incubated with 100 mM DTT for 20 min at 42 °C. A lithium acetate-based transformation mix was added, together with 1 μg of purified plasmid, and cells were transformed at 42 °C for 40 minutes. The transformed cells were selected on plates containing 400 mg/L Hygromycin B (Sigma-Aldrich, Espoo, Finland). Successful mating type change was determined by PCR as described below. Colonies from selection plates were replated three times onto YPD agar plates to encourage plasmid loss, after which they were stored at − 80 °C.

### PCR to confirm mating type change and hybridizations

The mating type locus was amplified with PCR using the previously published primers: MAT-R (AGTCACATCAAGATCGTTTATGG), MATa-F (ACTCCACTTCAAGTAAGAGTTTG) and MATα-F (GCACGGAATATGGGACTACTTCG) (Huxley et al. 1990). These primers amplify a 404-bp fragment for *MAT*α, and a 544-bp fragment for *MAT***a**. In addition, the presence of HMLα (new primers designed) and HMR**a** (Ota et al. 2018), was tested using the following primers: HMLα-F (GAATGGCACGCGGACAAAAT), HMLα-R (TGGAACACAGAAAAGAGCAGTG), HMR**a**-F (GTTGCAAAGAAATGTGGCATTACTCCA), HMR**a**-R (AGCTTTCTCTAACTTCGTTGACAAA). Interdelta fingerprints were produced using delta12 and delta21 primers from Legras and Karst (2003). PCR reactions were carried out with Phusion High-Fidelity PCR Master Mix with HF Buffer (Thermo Scientific, Vantaa, Finland) and primer concentrations of 0.5 μM. PCR products were separated and visualized on 1.0% agarose gels or on an Agilent ZAG DNA Analyzer capillary electrophoresis device.

### Hybridizations and selection of meiotic segregants

Hybridizations between mating-competent variants were attempted by placing cells of both parent strains, with opposite mating types, adjacent to each other on a YPD agar plate with the aid of a MSM400 dissection microscope (Singer Instruments, UK). Plates were incubated at 25 °C for up to 5 days, after which any emerging colonies were replated twice on fresh YPD plates to ensure single colony isolates. PCR of the mating type locus and interdelta fingerprints were used to confirm successful hybridization.

Selected hybrids were transferred to 1% potassium acetate agar for sporulation. After 7 days of incubation at 25 °C, ascospores were digested (using Zymolyase 100T) and dissected on YPD agar using the MSM400 dissection microscope.

### IRC7 copy number estimation by qPCR

The relative copy numbers of the *IRC7* gene in selected strains was estimated with quantitative PCR of genomic DNA. Primers PF6 and PR7 from Roncoroni et al. (2011) were used for *IRC7*. Copy numbers were normalized to that of *ALG9* and *UBC6* (primers listed in Krogerus et al. (2019)). The efficiencies (E) of the qPCR assays (ranging from 1.9 to 1.94) for each primer pair were calculated using the formula 10(−1/m), where m is the slope of the line of the threshold cycle (CT)-versus-log dilution plot of the DNA template (8 pg to 8 ng input DNA) (Pfaffl 2001). The qPCR reactions were prepared with PerfeCTa SYBR® Green SuperMix (QuantaBio, Beverly, MA, USA) and 0.3 μM of the primers. The qPCR reactions were performed on a LightCycler® 480 II instrument (Roche Diagnostics, Basel, Switzerland) in four technical replicates on 1 ng template DNA. The following programme was used: pre-incubation (95 °C for 3 min), amplification cycle repeated 45 times (95 °C for 15 s, 60 °C for 30 s, 72 °C for 20 s with a single fluorescence measurement), melting curve programme (65–97 °C with continuous fluorescence measurement), and finally a cooling step to 40 °C. The copy numbers of *IRC7* relative to *ALG9* and *UBC6* were calculated using the Pfaffl method (Pfaffl 2001).

### DNA content by flow cytometry

Ploidy of selected strains was measured using SYTOX Green staining and flow cytometry as described previously (Krogerus et al. 2017).

### Whole-genome sequencing and analysis

For analysis of the parent strains, sequencing reads were first obtained from NCBI-SRA (accession numbers in Supplementary Table S1). Reads were trimmed and filtered with fastp using default settings (version 0.20.1; Chen et al., 2018). Trimmed reads were aligned to a *S. cerevisiae* S288C reference genome (Engel et al. 2014) using BWA-MEM (Li and Durbin 2009), and alignments were sorted and duplicates were marked with sambamba (version 0.7.1; Tarasov et al., 2015). Variants were jointly called in all strains using FreeBayes (version 1.32; Garrison and Marth, 2012). Variant calling used the following settings: --min-base-quality 30 --min-mapping-quality 30 --min-alternate-fraction 0.25 --min-repeat-entropy 0.5 --use-best-n-alleles 70 -p 2. The resulting VCF file was filtered to remove variants with a quality score less than 1000 and with a sequencing depth below 10 per sample using BCFtools (Li 2011). Variants were annotated with SnpEff (Cingolani et al. 2012).

For phylogenetic analysis, the variants were filtered to retain only single nucleotide polymorphisms and remove sites with a minor allele frequency less than 5%. The filtered SNP matrix was converted to PHYLIP format (https://github.com/edgardomortiz/vcf2phylip). A random allele was selected for heterozygous sites. A maximum likelihood phylogenetic tree was generated using IQ-TREE (version 2.0.3; Nguyen et al. 2015) run with the ‘GTR+G4’ model and 1000 bootstrap replicates (Minh et al. 2013).

Four hybrid strains were whole-genome sequenced at NovoGene (UK). DNA was extracted using the method described by Denis et al. (2018). Sequencing was carried out on an Illumina NovaSeq 6000 instrument. The 150bp paired-end reads have been submitted to NCBI-SRA under BioProject number PRJNA740182. Analysis of the hybrid strains was carried out essentially as described above. Sequencing coverage was estimated with mosdepth (version 0.2.6; Pedersen and Quinlan 2018). Chromosome copy numbers were estimated based on distribution of alternate allele frequencies, ploidy as measured by flow cytometry, and sequencing coverage.

### Wort fermentations

Lab-scale wort fermentations were first carried out in triplicate to screen the yeast hybrids in order to identify top-performing strains that were able to rapidly attenuate wort sugars. To do this, overnight cultures of the yeast hybrids were set up by inoculating single colonies in 10 mL wort. These were then incubated at 25 °C with shaking (120 rpm) for 24 hours. The optical density (OD_600_) of the overnight cultures was measured and the cultures were diluted into 400 mL 10 °P wort (made from pale barley malt and 2.4 g L^-1^ of Cascade hops) in 500 mL glass bottles to a starting OD600 value of 0.3 as previously described by Mertens et al. (2015). The bottles were fitted with airlocks and were incubated at 25 °C. Specific gravity readings of the fermenting wort were taken daily for 7 days using the DMA 35 handheld density meter (Anton Paar GmbH, Austria).

2L-scale wort fermentations were carried out in 3-L cylindroconical stainless steel fermenting vessels, containing 2 L of 15 °P wort. Yeast was propagated in autoclaved wort. The 15 °P wort (70.5 g maltose, 21 g maltotriose, 19 g glucose, and 4.6 g fructose per liter) was produced at the VTT Pilot Brewery from barley malt and contained 2.5 g L^−1^ each of Cascade and Perle hops added to the whirlpool. The wort was oxygenated to 10 mg L^−1^ prior to pitching (Oxygen Indicator Model 26073 and Sensor 21158; Orbisphere Laboratories, Switzerland). Yeast was inoculated at a rate of 15 × 10^6^ viable cells mL^−1^, together with 2.5 g L^−1^ each of Cascade and Perle hops (dry hopping). The fermentations were carried out in triplicate at 20 °C until no change in alcohol level was observed for 24 h or for a maximum of 9 days.

Wort samples were drawn regularly from the fermentation vessels aseptically and placed directly on ice, after which the yeast was separated from the fermenting wort by centrifugation (9000×g, 10 min, 1 °C).

### Beer chemical analysis

The specific gravity, alcohol level (% v/v), and pH of samples were determined from the centrifuged and degassed fermentation samples using an Anton Paar Density Metre DMA 5000 M with Alcolyzer Beer ME and pH ME modules (Anton Paar GmbH, Austria).

Concentrations of fermentable sugars (glucose, fructose, maltose, and maltotriose) and ethanol were measured by HPLC using a Waters 2695 Separation Module and Waters System Interphase Module liquid chromatograph coupled with a Waters 2414 differential refractometer (Waters Co., Milford, MA, USA). An Aminex HPX-87H Organic Acid Analysis Column (300 × 7.8 mm; Bio-Rad, USA) was equilibrated with 5 mM H_2_SO_4_ (Titrisol, Merck, Germany) in water at 55 °C, and samples were eluted with 5 mM H_2_SO_4_ in water at a 0.3 mL min^−1^ flow rate.

Higher alcohols and esters were determined by headspace gas chromatography with flame ionization detector (HS-GC-FID) analysis. Four-milliliter samples were filtered (0.45 μm) and incubated at 60 °C for 30 min, and then 1 mL of gas phase was injected (split mode; 225 °C; split flow of 30 mL min^−1^) into a gas chromatograph equipped with an FID detector and headspace autosampler (Agilent 7890 Series; Palo Alto, CA, USA). Analytes were separated on a HP-5 capillary column (50 m × 320 μm × 1.05 μm column; Agilent, USA). The carrier gas was helium (constant flow of 1.4 mL min^−1^). The temperature program was 50 °C for 3 min, 10 °C min^−1^ to 100 °C, 5 °C min^−1^ to 140 °C, 15 °C min^−1^ to 260 °C and then isothermal for 1 min. Compounds were identified by comparison with authentic standards and were quantified using standard curves. 1-Butanol was used as internal standard.

4-Vinyl guaiacol was analyzed using HPLC based on methods described by Coghe et al. (2004) and McMurrough et al. (1996). The chromatography was carried out using a Waters Alliance HPLC system consisting of a Waters e2695 Separations Module equipped with a XTerra® MS C18 column (5 µm, 4.6 × 150 mm) and a Waters 2996 Photodiode Array Detector. The mobile phase consisted of H_2_O/CH_3_OH/H_3_PO_4_ (64:35:1, v/v) and flow rate was 0.5 mL min^−1^. The diode array detector was used at 190–400 nm. 4-Vinyl guaiacol was quantified at 260 nm using standard curves of the pure compound (0.3–10 mg L^−1^).

The volatile thiols 4-mercapto-4-methyl-2-pentanone (4MMP), 3-mercapto-1-hexanol (3MH), and 3-mercaptohexylacetate (3MHA) were determined using the method described by Dennenlöhr et al. (2020). In this method thiols are extracted and derivatized by headspace solid-phase microextraction (HS-SPME) with on-fiber derivatization (OFD) using 2,3,4,5,6-pentafluorobenzyl bromide (PFBBr). Resulting PFBBr-thioesters are then separated and analysed using gas chromatography tandem mass spectrometry (GC-MS/MS). The instrumental setup, parameters of sample preparation, GC-MS/MS analysis, calibration, and quantification were in full accordance to Dennenlöhr et al. (2020). Each sample was analysed in duplicate.

### Data visualization and analysis

Data and statistical analyses were performed with R (http://www.r-project.org/). The phylogenetic tree was produced using the ‘ggtree’ package (Yu et al. 2017). Flow cytometry data was analysed with ‘flowCore’ (Hahne et al. 2009) and ‘mixtools’ (Benaglia et al. 2009) packages. Scatter and box plots were produced with the ‘ggpubr’ package (Kassambara 2020). Variants along the genome were visualized in R using the ‘karyoploter’ package (Gel and Serra 2017).

## Results

### Identifying suitable parent strains for improving β-lyase activity

As the aim of the applied part of this study was to obtain brewing yeast strains with improved β-lyase activity, we first performed a phenotypic and genetic pre-screening step to identify suitable parent strains to use for the CRISPR-mediated hybridizations. A set of thirty-eight *Saccharomyces cerevisiae* strains were included in the screening (Supplementary Table S1). Thirty-seven of these were brewing strains from Escarpment Laboratories, while the final strain, YJM1400 (or SACE_YCM), was selected from the 1,011 yeast genomes study (Peter et al. 2018). This strain was included here, as we predicted it to have a high β-lyase activity based on its *IRC7* sequence and copy number. The main β-lyase enzyme in *S. cerevisiae* is encoded by the *IRC7* gene (Roncoroni et al. 2011; Ruiz et al. 2021). YJM1400 not only carries the more active full-length allele of *IRC7* (Roncoroni et al. 2011), but also lacks any of the widespread inactivating mutations that have recently been identified (Cordente et al. 2019), and appears to be one of the few strains in the 1,011 yeast genomes study with enhanced *IRC7* copy number.

Using whole-genome sequence data, we first queried the presence of inactivating mutations (Cordente et al. 2019) in *IRC7* among the 38 strains. The long allele of *IRC7* was present in most strains, while the Thr185Ala missense mutation was common among the brewing strains (Figure 1). The Thr185Ala mutation could be found, for example, among all strains in the ‘United Kingdom’ sub-clade, where it was often homozygous. Other inactivating mutations were also frequent among the tested strains, including Lys43Arg, Tyr56*, His197Gln and Val348Leu (Cordente et al. 2019; Curtin et al. 2020). Only a handful of strains lacked any of the known inactivating mutations, and these were thus predicted to have a higher β-lyase activity. In addition to *IRC7*, we queried for the presence of loss-of-function mutations in *URE2*, which encodes a regulatory protein involved in nitrogen catabolite repression, which reduces *IRC7* expression (Thibon et al. 2008; Dufour et al. 2013). A group of five brewing strains were found to contain a heterozygous nonsense mutation in *URE2* (Figure 1). Presence of inactivating mutations in *URE2* have been shown to increase volatile thiol release during wine fermentations (Dufour et al. 2013).

**Figure 1.**
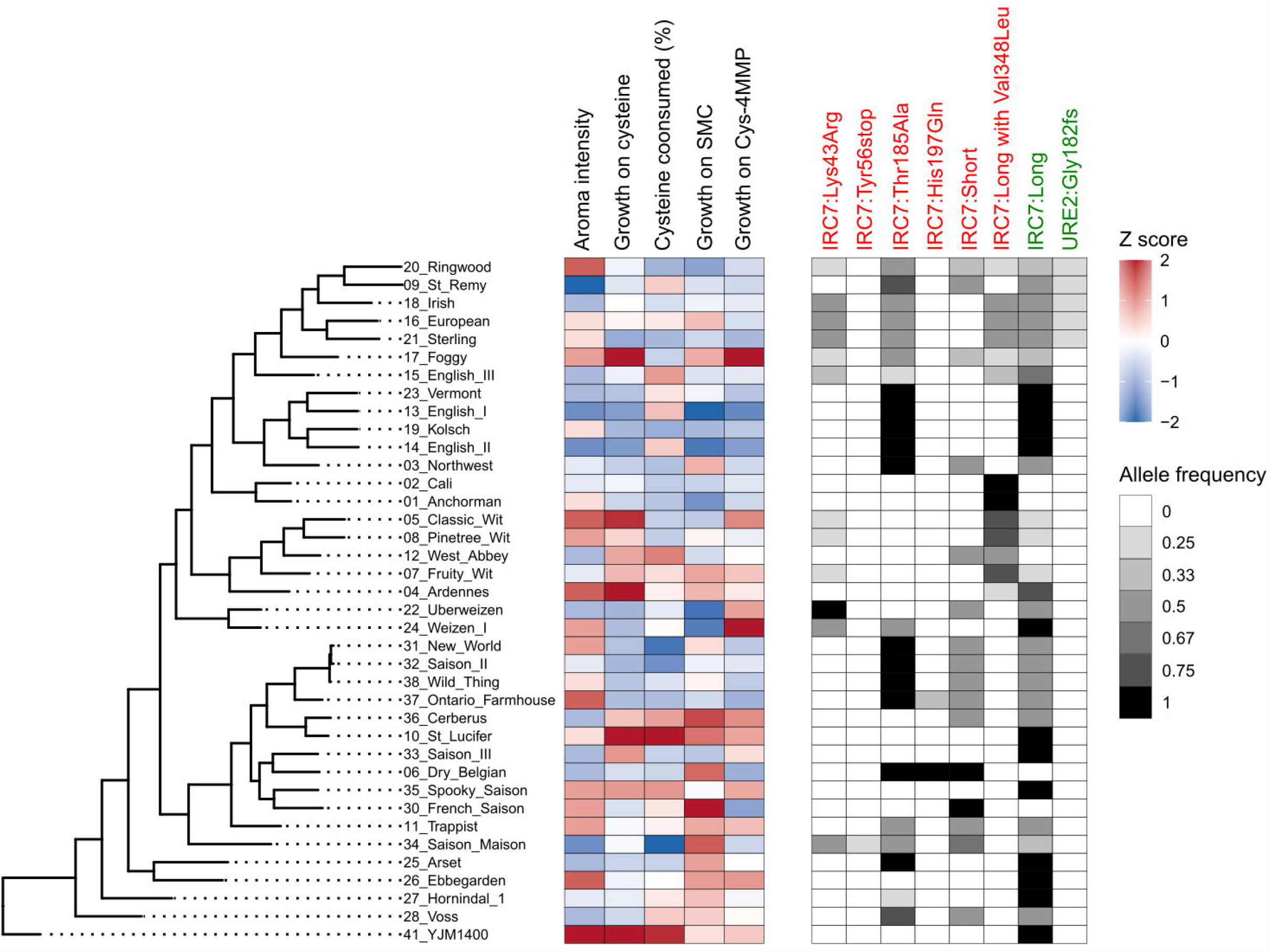
Genetic and phenotypic screening of β-lyase activity in the 38 *Saccharomyces cerevisiae* included in the study. Strains are ordered based on phylogenetic relationship (maximum likelihood phylogenetic tree based on SNPs at 114700 sites, rooted with *S. cerevisiae* YJM1400 as outgroup). The phenotypic heatmap is colored blue to red based on Z-scores. The genotypic heatmap is colored from white to black based on allele frequency of the different mutations. The mutations that are colored red have been shown to decrease β-lyase activity, while the mutations colored green have been shown to increase β-lyase activity.

In addition to the genetic pre-screening, the β-lyase activity of the strains was estimated by testing their growth on various cysteine-conjugates as sole nitrogen source, and by scoring aroma intensity after fermentation in wort supplemented with Cys-4MMP. In general, there was good agreement between the phenotype and genotype, as the best performing strains (e.g. YJM1400, St. Lucifer, Spooky Saison, Ardennes, and Ebbegarden) were those without or with rare heterozygous inactivating mutations. For strains containing a homozygous long allele of *IRC7*, significantly higher growth on cysteine and aroma intensity from wort supplemented with Cys-4MMP was observed in strains without any inactivating mutations compared to those with homozygous inactivating mutations (Supplementary Figure S1). Between phenotypes, moderate positive correlation was also observed between many of the measured phenotypes (Supplementary Table S2).

Based on these pre-screenings, we selected eight candidate parent strains for the hybridization trials (Table 1). Four of these were selected based on high predicted β-lyase activity in the pre-screenings, and they included YJM1400, St. Lucifer, Ardennes and Classic Wit. As the end goal was to develop yeast strains suitable for the production of IPA-style beers, where phenolic off-flavours are unwanted, the remaining four strains were selected among the pool of POF-strains. These strains were Cerberus, Ebbegarden, Foggy London and Sterling. Next, we attempted to generate mating-competent variants of these eight strains using the CRISPR/Cas9 system.

**Table 1.** Colonies appearing on selection plates after transformation by Cas9 plasmid targeting *MAT***a** or MATα.

| Strain | *MATa* plasmid | *MATα* plasmid |
| --- | --- | --- |
| Ardennes | 11 | 18 |
| Classic Wit | 5 | 5 |
| St Lucifer | 4 | 7 |
| Foggy | 4 | 1 |
| Sterling | 28 | 10 |
| Ebbegarden | 8 | 13 |
| Cerberus | 4 | 3 |
| YJM1400 | 63 | 133 |

### Generating mating-competent variants for hybridization

The eight parent strains that were selected based on pre-screenings were transformed with CRISPR/Cas9 plasmids containing protospacer sequences targeting either *MAT***a** and *MAT*α using optimized stationary phase transformation (Tripp et al. 2013). Transformation efficiencies varied broadly, with between 1 and 133 colonies emerging on the selection plates (400 mg hygromycin / mL) from the transformation of 1.5 mL saturated overnight culture (Table 1). Colonies were obtained for all 16 combinations (eight strains with two plasmids). Up to six colonies from each strain and plasmid were transferred to fresh selection plates, after which DNA was extracted and PCR was used to confirm successful mating-type change (Supplementary Figure S2). Out of the 80 colonies that were tested, mating-type change from *MAT***a**/*MAT*α to either *MAT***a** or *MAT*α had successfully occurred in 73.

Next, these 73 transformants were replated twice on non-selective media (YPD without hygromycin) to encourage loss of the CRISPR/Cas9 plasmid. Plasmid loss was confirmed in 66 transformants by lack of growth when replated back to selection plates. As all eight parent strains are homothallic, we were unsure if the mating-type would remain stable after loss of the CRISPR/Cas9 plasmid. In wild-type homothallic strains, mating-type change would occur at cell division following the repair of a double-stranded break (DSB) created by the HO endonuclease at the mating type locus. We retested the mating-type of all 66 transformants lacking the CRISPR/Cas9 plasmid by PCR, and all strains still exhibited a single mating-type. Stable *MAT***a** and *MAT*α variants were successfully obtained for all eight parent strains. The CRISPR-based mating-type switching process developed by Xie et al. (Xie et al. 2018) therefore appears to generate stable mating-competent variants even from homothallic industrial strains.

As the protospacer sequences used to target *MAT***a** and *MAT*α are also present in the silent mating-type cassettes *HMR***a** and *HML*α on either end of chromosome III, we hypothesized that the stable mating-type in the transformed homothallic strains are a result of simultaneous deletion of the respective silent mating-type cassettes. To test this, we performed PCR on wild-type and transformed strains using primers designed to amplify *HMR***a** and *HML*α. Wild-type strains yielded products with both primer pairs, while transformants only yielded single products, indicating that *HMR***a** and *HML*α are indeed deleted during the mating-type switching process (Supplementary Figure S3).

### Construction of hybrids

Following the successful isolation of stable mating-competent variants of the eight selected parent strains, we proceeded with hybridization attempts (Figure 2A). From these strains, we attempted 21 crosses in total. As the end goal was to obtain a yeast strain lacking the POF phenotype, each cross involved at least one POF-parent. Hybridizations were attempted by placing cells of both parent strains adjacent to one another on a YPD agar plate using a Singer MSM400 dissection microscope. 16 pairs per cross were placed together. Of the 21 attempted crosses, 18 successfully yielded hybrids (for a total of 63 hybrids; Supplementary Table S3). Hybridization frequency varied considerably between the successful crosses, ranging from 6.3 to 63% (median 18.8%, average 25%). Hybridization was confirmed by checking both for heterozygosity at the *MAT* locus using PCR (as both parent strains showed stable single mating types) and by producing interdelta fingerprints using PCR and capillary electrophoresis (examples in Figure 2B and C). In the interdelta fingerprints, successful hybrids produced bands of both parent strains. Flow cytometry and DNA staining with SYTOX Green of selected hybrids also revealed that ploidy of the hybrid strains had increased to levels above both parent strains (Table 2). Crossing of the tetraploid strain 21 (Sterling) and diploid strain 10 (St. Lucifer), for example, resulted in a hexaploid hybrid (Figure 2D). Indeed, of the seven hybrids of which the ploidy was measured, six appeared to be approximately hexaploid.

**Table 2.** Estimated ploidy of selected parent strains, hybrids and spore clones as measured by SYTOX Green-staining and flow cytometry.

| Strain | Type | Ploidy |
| --- | --- | --- |
| 10. St. Lucifer | Parent strain | 1.9 ± 0.09 |
| 17. Foggy | Parent strain | 4.1 ± 2.64 |
| 21. Sterling | Parent strain | 3.8 ± 0.18 |
| 26. Ebbegarden | Parent strain | 3.9 ± 0.13 |
| 36. Cerberus | Parent strain | 2.0 ± 0.14 |
| 41. YJM1400 | Parent strain | 2.1 ± 0.11 |
| 21 x 10 D4 | F1 hybrid | 6.0 ± 0.86 |
| 21 x 41 A2 | F1 hybrid | 6.0 ± 1.29 |
| 26 x 10 D1 | F1 hybrid | 5.5 ± 0.49 |
| 26 x 17 D3 | F1 hybrid | 5.7 ± 1.12 |
| 26 x 41 A3 | F1 hybrid | 5.9 ± 0.74 |
| 36 x 26 D3 | F1 hybrid | 5.9 ± 0.93 |
| 36 x 41 A2 | F1 hybrid | 4.0 ± 0.16 |
| 21 x 10 D4 C3 | F1 spore clone | 2.8 ± 0.13 |
| 21 x 41 A2 A3 | F1 spore clone | 2.8 ± 0.11 |
| 21 x 41 A2 B3 | F1 spore clone | 3.4 ± 0.65 |
| 21 x 41 A2 D1 | F1 spore clone | 2.9 ± 0.16 |
| 21 x 41 A2 E1 | F1 spore clone | 2.6 ± 0.12 |
| 26 x 10 D1 A2 | F1 spore clone | 5.0 ± 0.84 |
| 26 x 41 A3 B3 | F1 spore clone | 3.0 ± 0.21 |
| 36 x 41 A2 A3 | F1 spore clone | 2.1 ± 0.17 |

**Figure 2.**
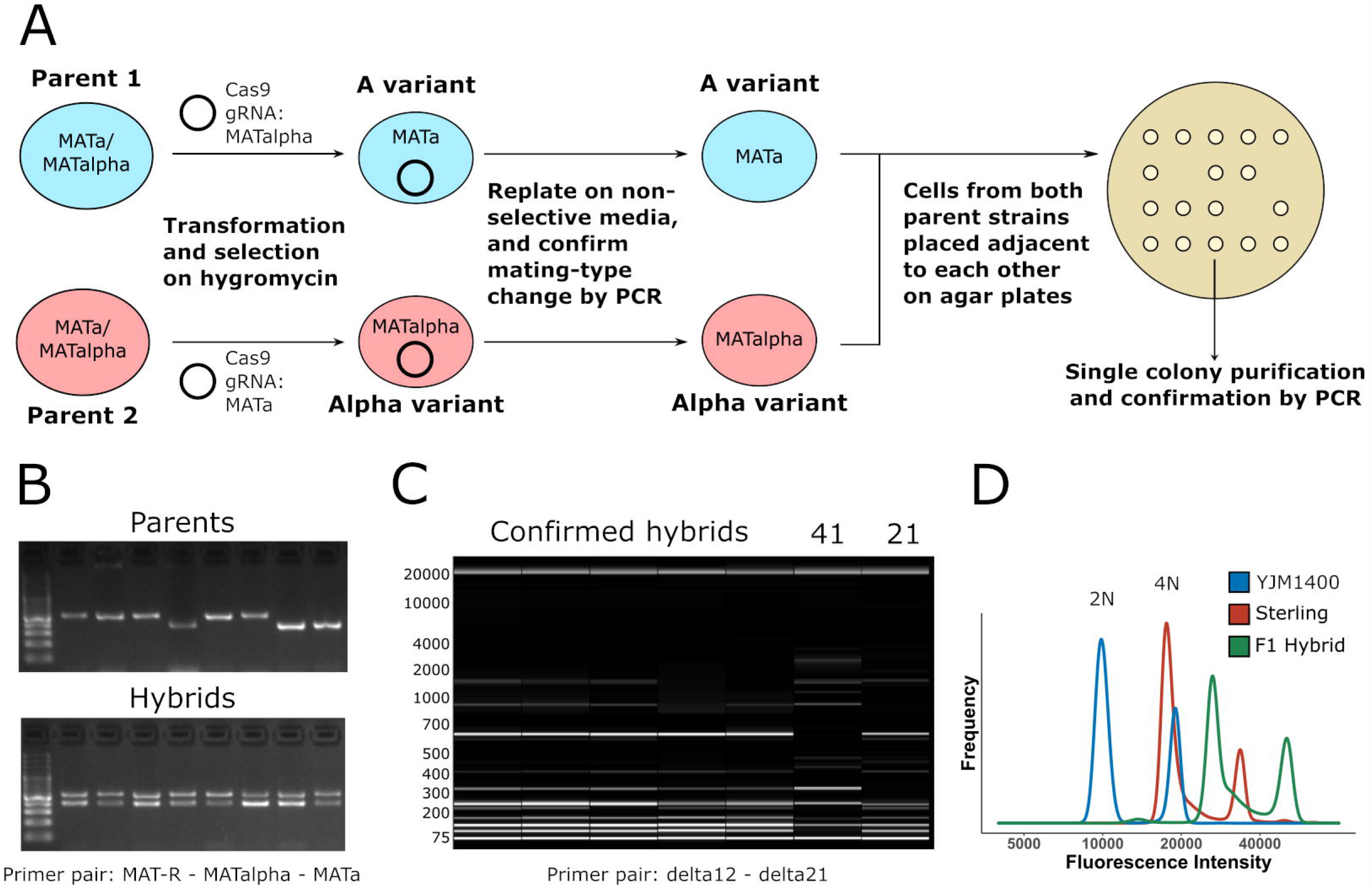
Overview of hybrid construction and confirmation. (**A**) Scheme of how parent strains were converted to mating-competent variants, which were then mated to form hybrids. (**B**) Mating type PCR to confirm hybridization. Parents produced a single band for either *MAT***a** or *MAT*α, while hybrids produced both bands. (**C**) Interdelta fingerprints to confirm hybridization. Hybrids produce fingerprints containing all the bands of the parent strains. (**D**) Flow cytometry and SYTOX Green staining reveal an increased ploidy of the F1 hybrid formed between Sterling and YJM1400 compared to the parent strains.

After hybrids were successfully constructed and confirmed, we still attempted to remove the POF phenotype from hybrid combinations involving a POF+ parent strain through meiotic segregation. Hybrids from five crosses were spread on potassium acetate agar for sporulation. Prior to sporulation, all successful hybrids from these crosses were screened for β-lyase activity by testing growth on cysteine as the sole nitrogen source. The best performing hybrid from each cross was chosen for sporulation. All five hybrids sporulated efficiently and formed viable spores, with spore viability ranging from 39% to 69%. A total of 47 spore clones were obtained. The ploidy of selected spore clones was measured with flow cytometry, and it had halved compared to the F1 hybrid in most cases (Table 2).

### Screening of constructed hybrids reveals heterosis

The spore clones, along with selected F1 hybrids and parent strains were first screened for various relevant traits in microplate format. These included efficient fermentation of wort, lack of phenolic off-flavour production, and ability to grow on cysteine as a sole nitrogen source (as an indicator of β-lyase activity). Considerable variation was observed among the screened traits in the 60 strains (Supplementary Figure S4). We decided to focus on the Sterling × YJM1400 hybrid (21 × 41 A2) and derived spore clones in more detail. In regards to the ability to grow on and consume 15 mM cysteine as a sole nitrogen source, we observed mid-parent heterosis in the F1 hybrid and derived spore clones (Figure 3A and B). Numerous spore clones outperformed the F1 hybrid. We also measured *IRC7* copy number (normalized to the copy numbers of *ALG9* and *UBC6* that were chosen as reference genes) by quantitative PCR, and observed a moderately strong positive correlation between *IRC7* copy numbers and the measured phenotypes (Figure 3C and D).

**Figure 3.**
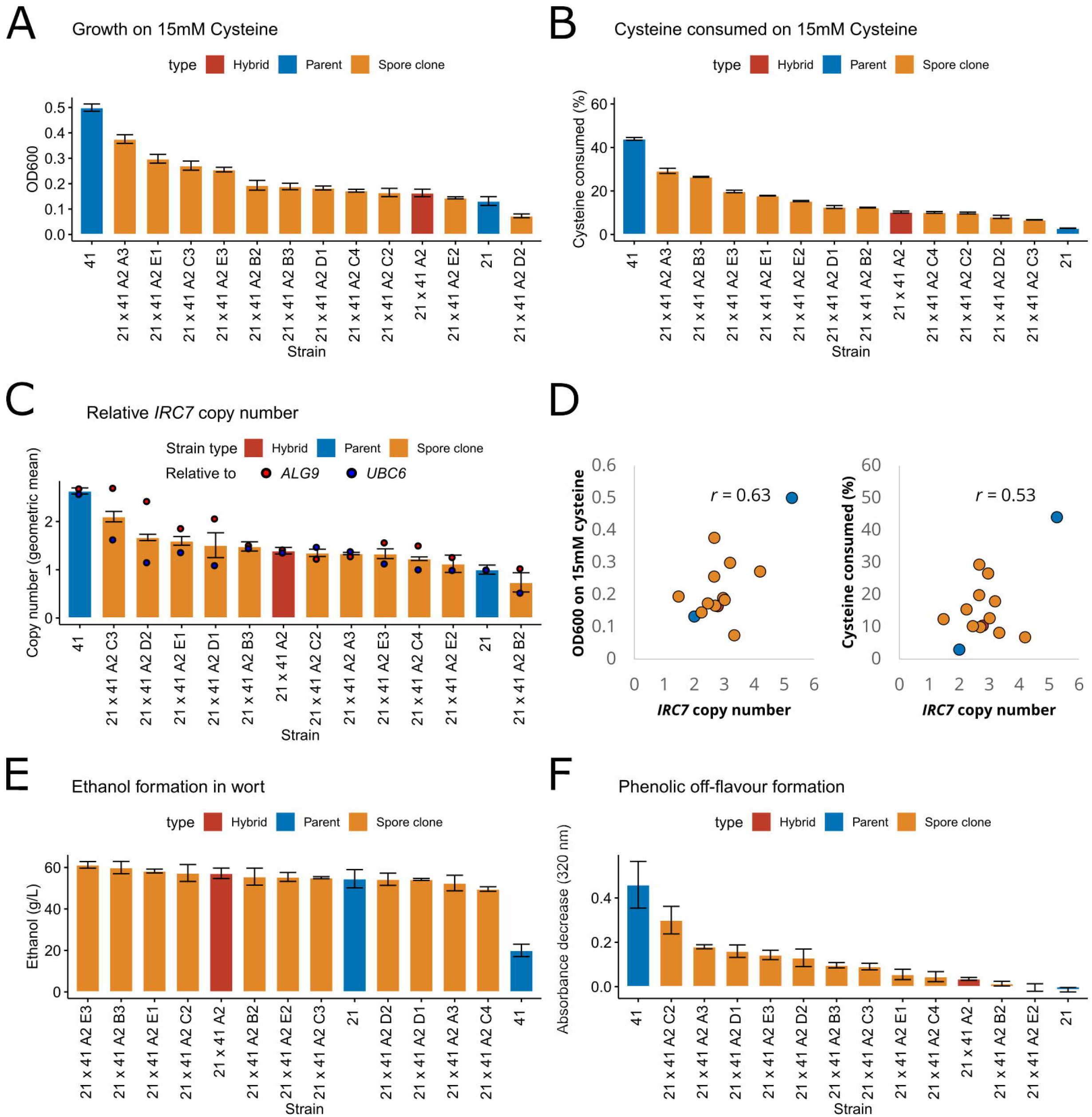
Phenotypic screening of Sterling × YJM1400 hybrid and eleven spore clones. (**A**) The OD600 reached when grown on 15mM cysteine as sole nitrogen source. (**B**) The amount of cysteine consumed during the cultivations on 15mM cysteine as sole nitrogen source. (**C**) The relative *IRC7* copy number normalized to *ALG9* and *UBC6*, as determined by quantitative PCR. (**D**) The correlation between *IRC7* copy number and growth on 15mM cysteine as sole nitrogen source. (**E**) The amount of ethanol (g/L) produced from 15 °P wort in microplate fermentations. (**F**) The decrease in absorbance at 320 nm after cultivations in 100 mg/L ferulic acid. Assays were done in triplicate, and error bars represent standard deviation. 21: Sterling. 41: YJM1400.

The F1 hybrid and spore clones, along with the ale parent Sterling, fermented wort efficiently, with measured ethanol levels ranging from around 50-60 g/L (Figure 3E). The wild parent YJM1400 only reached 20 g ethanol/L, indicating it was unable to ferment maltose and maltotriose from the wort. Strains were grown in the presence of ferulic acid to test phenolic off-flavour (POF) formation. Greatest conversion of ferulic acid to 4-vinylguaiacol was as expected observed for the wild parent YJM1400 (Figure 3F). Interestingly, barely any drop in absorbance was observed with the F1 hybrid, despite it containing functional alleles of *PAD1* and *FDC1* from YJM1400. Similarly to the other traits, considerable variation was observed among the spore clones. *PAD1* and *FDC1* were Sanger-sequenced in the fourteen strains (two parents, one F1 hybrid, and eleven spore clones) to clarify the results of the POF assay (Supplementary Figure S5). Homozygous loss-of-function (LOF) mutations in *PAD1* and *FDC1* were observed in the ale parent Sterling, as well as two out of eleven spore clones (A2 B2 and A2 C4). This ratio (0.18) corresponds well to the predicted ratio of spore clones being homozygous for the LOF mutations, assuming the spore clones are triploid and the hexaploid hybrid has four LOF alleles and two functional alleles (*C*(4,3) / *C*(6,3) = 0.2).

After high-throughput screening, a total of seven hybrids and eight spore clones, along with the six parent strains, were selected for 400mL-scale wort fermentations (Figure 4A). Best-parent heterosis was observed in regards to fermentation rate, as F1 hybrids reached the mid-point of fermentation significantly faster than the parent strains (Figure 4B). F1 hybrids also reached, on average, a higher attenuation level, but the difference to the parent strains was not significant (Figure 4C). The spore clones appeared to perform on average slightly worse than the F1 hybrids in regards to fermentation rate and final attenuation, however, the difference was not significant (*p* > 0.05).

**Figure 4.**
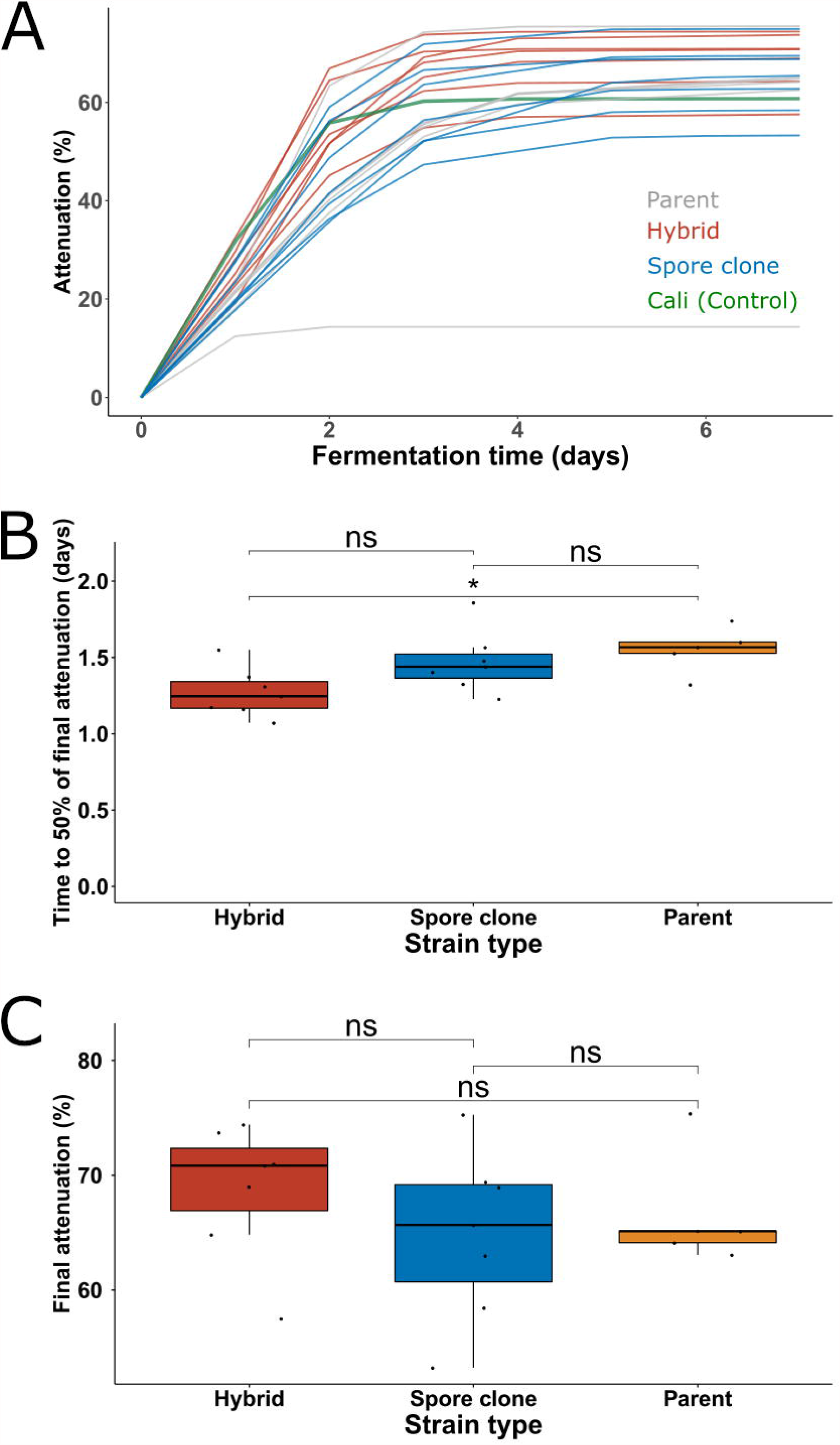
Screening of parent, hybrid and spore clone strains in 400mL wort fermentations. (**A**) The apparent attenuation (%) during fermentations. Curves are colored according to strain type. Cali Ale was included as a control. (**B**) The time taken to reach 50% of the final attenuation in the different strain groups. (**C**) The final attenuation reached in the different strain groups. Groups were compared with the Wilcoxon signed-rank test, and an asterisk (*) indicated *p* < 0.05. ns: not significant. Fermentations were performed in triplicate.

### Confirmation of enhanced phenotype in 2L-scale wort fermentations

Two F1 hybrids and two derived spore clones were selected for 2L-scale wort fermentations and more detailed phenotyping. Both hybrids involved the wild parent YJM1400, with the other parent being one of two brewing strains, Sterling or Ebbegarden. As was already observed during the smaller scale wort fermentations, the F1 hybrids exhibited best-parent heterosis in regards to fermentation rate (Figure 5A). The Sterling × YJM1400 hybrid 21 × 41 A2, for example, had reached 3.9% ABV after 23 hours compared to 2.4% in Sterling (*p =* 0.001). The wild parent YJM1400 was unable to utilize the maltose and maltotriose in the wort, and only reached 1.4% ABV. The fermentation profile of the Sterling × YJM1400 spore clone 21 × 41 A2 E1 was identical to the Sterling parent, while the Ebbegarden × YJM1400 spore clone 26 × 41 A3 B3 fermented slower than the F1 hybrid or ale parent.

**Figure 5.**
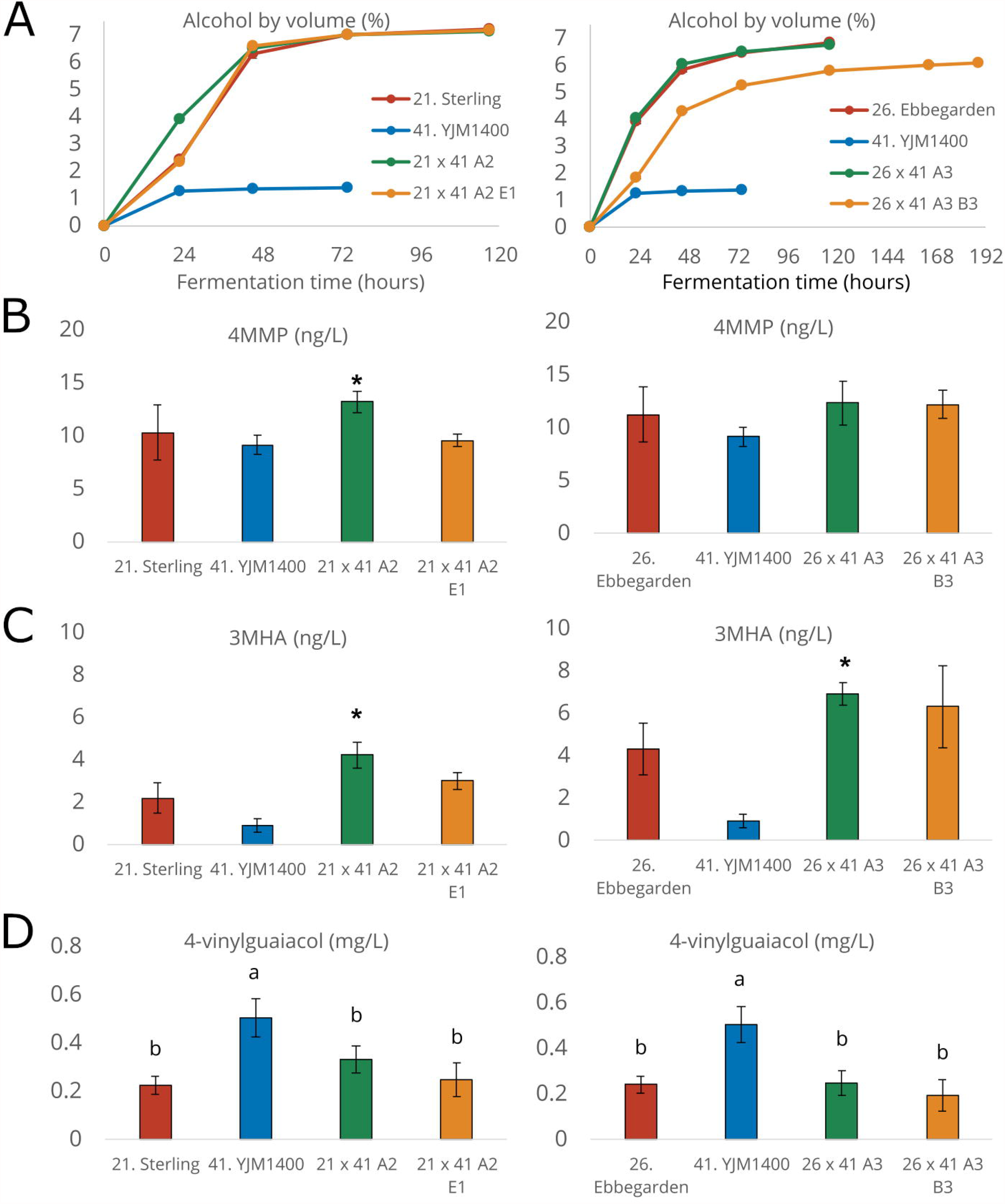
Fermentation performance and concentrations of thiols and 4-vinylguaicol in 2L-scale fermentations. (**A**) Alcohol by volume (%) during fermentations. Concentrations (ng/L) of (**B**) 4-mercapto-4-methyl-2-pentanone (4MMP), and (**C**) 3-mercaptohexylacetate (3MHA) in the beers. An asterisk (*) indicates a concentration significantly higher (*p* < 0.05) than both parent strains as determined by unpaired two-tailed t-test. (**D**) Concentrations of 4-vinylguaicol (mg/L) in the beers. Different letters indicate significant differences (*p* < 0.05) as determined by one-way ANOVA and Tukey’s post-hoc test. Fermentations were performed in triplicate.

The concentrations of 4MMP and 3MHA in the finished beers were also measured (Figure 5B and C). A significant increase in 4MMP was observed for the Sterling × YJM1400 hybrid 21 × 41 A2 compared to the parent strains (*p* < 0.05), while significant increases in 3MHA were observed for both the F1 hybrids. Concentrations of both 4MMP and 3MHA were above or around the flavour threshold (1 and 4 ng/L, respectively (Capone et al. 2018)) in the beers fermented with the hybrid strains, indicating a positive influence on flavour. Interestingly, despite the high apparent β-lyase activity in the YJM1400 strain, the beers made with this strain had low amounts of volatile thiols. It is possible that this is a result of the limited fermentation.

In addition to attempting to increase thiol formation, our goal was also to decrease 4-vinylguaiacol (4VG) formation in our hybrids. The F1 hybrids and spore clones produced lower levels of 4VG than the wild *S. cerevisiae* YJM1400 parent, which was the only strain that clearly produced levels above the flavour threshold of around 0.3 mg/L (Vanbeneden et al. 2008) (Figure 5D). 4VG concentrations of the hybrid beers were marginally higher than those measured in the beers fermented with the POF-parent strains (Sterling and Ebbegarden). Concentrations of yeast-derived esters were also enhanced in several of the beers produced with the hybrid strains (Figure 6A to C). The Ebbegarden × YJM1400 spore clone 26 × 41 A3 B3, in particular, produced higher levels of 3-methylbutyl acetate, ethyl hexanoate and ethyl octanoate compared to either parent. We also measured the flocculation potential of the strains (a desirable trait in brewing strains), and it remained as high as in the ale parent for three out of the four hybrid strains (Figure 6D).

**Figure 6.**
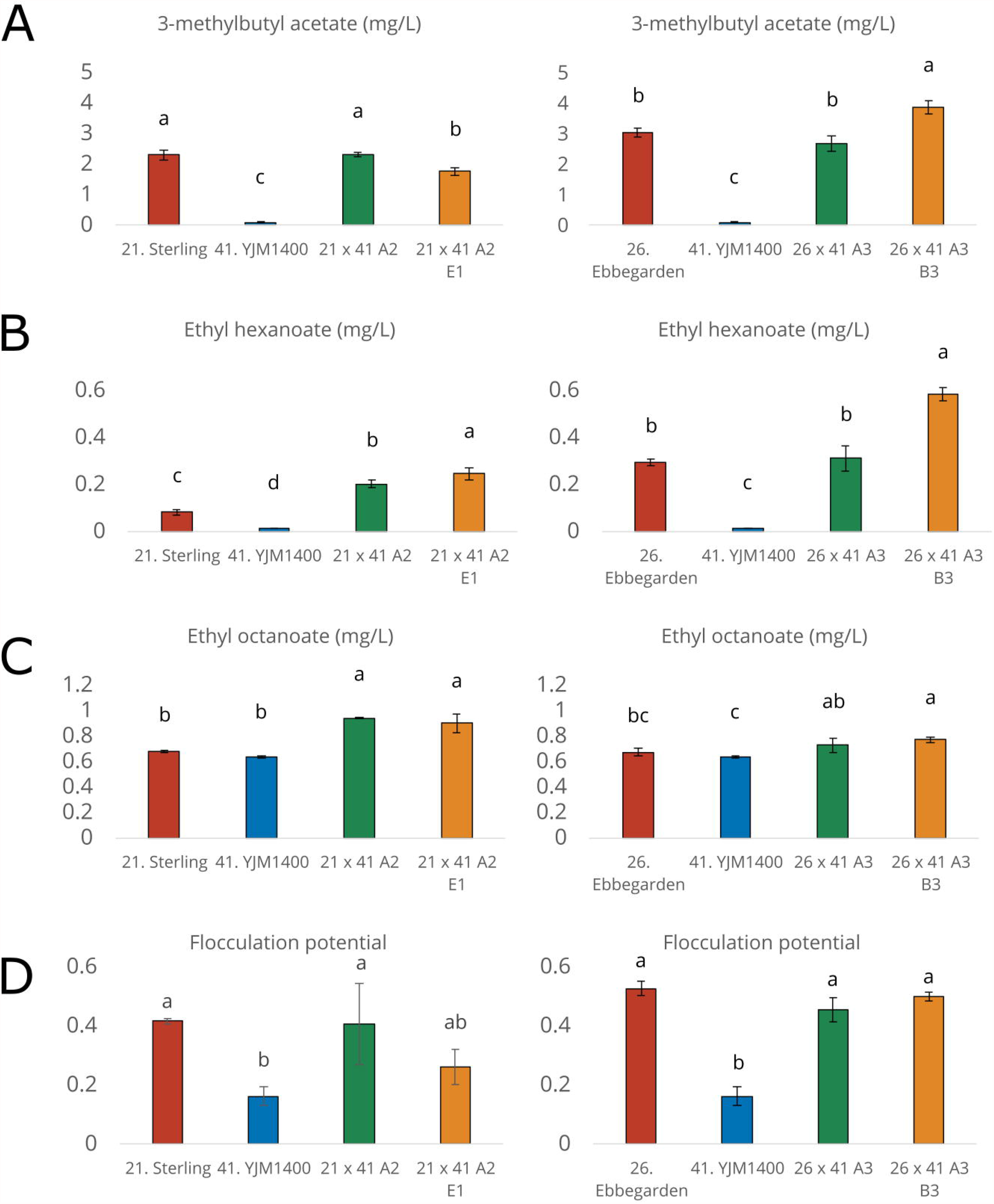
Ester concentrations and flocculation potential in 2L-scale fermentations Concentrations of (mg/L) of (**A**) 3-methylbutyl acetate, (**B)** ethyl hexanoate, and (**C**) ethyl octanoate in the beers. (**D**) Flocculation potential as determined by Helm’s test. Different letters indicate significant differences (*p* < 0.05) as determined by one-way ANOVA and Tukey’s post-hoc test. Fermentations were performed in triplicate.

### Whole-genome sequencing of the selected hybrid strains

The four hybrid strains that were studied in more detail above were whole-genome sequenced. The F1 hybrids were nearly euploid, having six copies of almost all chromosomes (Figure 7A). The spore clones had more variation in chromosome copy numbers, ranging from two to four. The F1 hybrids had high levels of heterozygosity, as over 100k heterozygous variants were identified in both hybrids (Figure 7B). Loss of heterozygosity (LOH) had occurred in the F1 spore clones, as the number of heterozygous variants decreased with approx. 20%. When the parent strains were compared to each other, a total of 50385 and 36642 variants unique to each parent were identified when Sterling and YJM1400 were compared, respectively. When Ebbegarden and YJM1400 were compared, 48157 and 32509 variants unique to each parent were identified, respectively. LOH had occurred for 1222 and 326 of the variants unique to Sterling and YJM1400, respectively, in the Sterling × YJM1400 spore clone 21 × 41 A2 E1, as they were now homozygous (Figure 7B and C). Similarly, 1038 and 1194 of the variants unique to Ebbegarden and YJM1400, respectively, were now homozygous in the Ebbegarden × YJM1400 spore clone 26 × 41 A3 B3. When plotted along the genome, these parent-specific homozygous sites were spread across the whole genome (Figure 7C).

**Figure 7.**
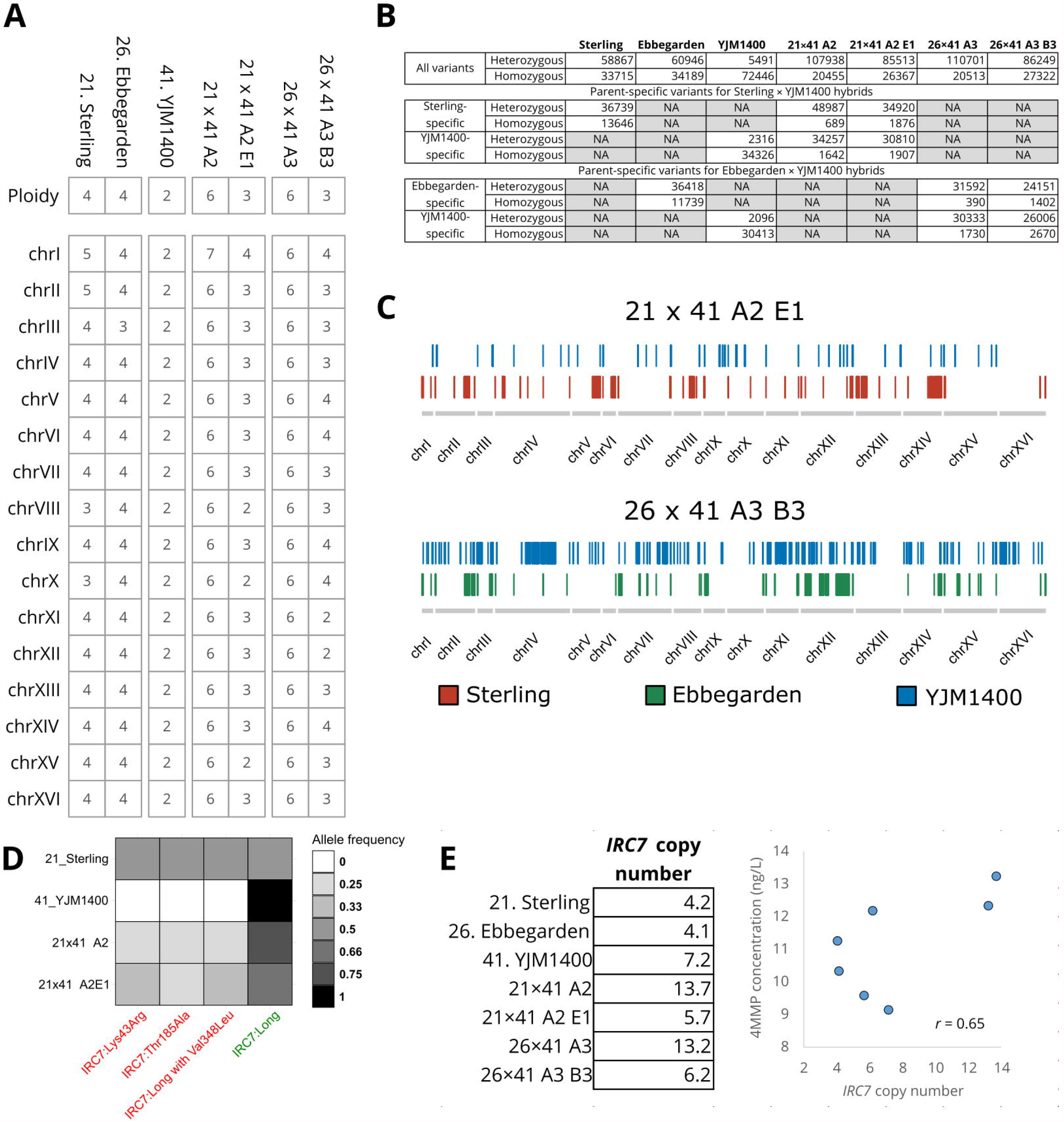
Whole-genome sequencing of selected hybrids and spore clones. (**A**) The estimated chromosome copy number and measured ploidy of the parent strains, hybrids and spore clones. (**B**) The amount of heterozygous and homozygous variants (compared to the *S. cerevisiae* S288C reference genome) detected in the strains. (**C**) Loss-of-heterozygosity regions where parent-specific mutations were homozygous in the spore clones (mutations were all heterozygous in the F1 hybrids). (**D**) Allele frequencies of the *IRC7* mutations in the Sterling × YJM1400 strains. (**E**) The estimated copy number of *IRC7* in the strains and correlation with beer 4MMP concentrations. Copy number was estimated based on median read coverage across *IRC7*, normalized to the read coverage across chromosome VI where the gene is located.

In regards to the β-lyase encoding *IRC7* gene, we saw both differential distribution of inactivating mutations and gene copy numbers among the parent and hybrid strains (Figure 7D and E). The Sterling parent strain contained three inactivating mutations with 50% allele frequency. These mutations were detected in the derived hybrids, but at a lower allele frequency (Figure 7D). None of the known inactivating mutations in *IRC7* (Figure 1) were observed in the Ebbegarden and YJM1400 parent strains, nor in their derived hybrids. *IRC7* copy numbers in the strains were estimated based on median coverage across the gene, normalized to the coverage across chromosome VI on which it is located. The copy numbers were as expected highest in the F1 hybrids, but decreased in the spore clones (Figure 7E). Nevertheless, *IRC7* copy numbers in the spore clones appeared higher than in the respective ale parent from which they were derived. A moderate positive correlation (*r* = 0.65) was observed between *IRC7* copy numbers and amount of 4MMP in the beers fermented with the strains.

## Discussion

Breeding with brewing yeast can be challenging, as most strains sporulate poorly or are unable to form viable spores (Gallone et al. 2016; De Chiara et al. 2020). Such strains can be bred with ‘rare mating’ (Gunge and Nakatomi 1972), but the approach is time-consuming and often not successful. Here, we set out to evaluate whether breeding of sterile industrial strains can be facilitated using CRISPR/Cas9-aided mating-type switching (Xie et al. 2018). Our results reveal that single mating type variants of industrial polyploid strains can indeed be readily generated and isolated. Interestingly, the mating type remained stable even after loss of the Cas9 plasmid, despite the strains being homothallic. Wild-type homothallic strains would, by action of the *HO*-coded endonuclease, switch mating type at cell division, and subsequently self-mate to reform a cell heterozygous at the mating type locus (Merlini et al. 2013). Here, no such mating type switching was observed, likely from the simultaneous loss of the silent mating type cassettes during the Cas9 transformations. This allows for the easy construction and maintenance of a library of mating-competent variants for large-scale breeding projects.

The single mating type variants readily mated with cells of opposite mating type, which allowed rapid construction of a large set of intraspecific hybrids. Numerous studies have demonstrated how breeding can be used combine and enhance traits from diverse strains (Steensels et al. 2014; Krogerus et al. 2015; Mertens et al. 2015; Krogerus et al. 2016). Hence, the approach used here can accelerate and simplify brewing yeast development through breeding, where hybrid construction would otherwise typically be the bottleneck. Here, we also observed heterosis for a number of traits in multiple hybrids, including fermentation rate and aroma formation. While not tested here, it is likely that the same approach, following modification of the protospacer sequences, could be applied to other *Saccharomyces* species as well to allow construction of interspecific hybrids. Furthermore, hybrids could also likely be retransformed to form mating-competent cells that could be bred with another parent strain. This would allow construction of multi-parent complex hybrids, such as those described by Peris et al. (2020).

Here, we aimed specifically at enhancing the β-lyase of selected brewing yeast strains. Volatile thiols have a central role in contributing fruity hop aroma in beer, and they typically are abundant in modern heavily-hopped IPA-style beers (Gros et al. 2012; Cibaka et al. 2017; Dennenlöhr et al. 2020; Bonnaffoux et al. 2021). As the vast majority of all thiols in hops are cysteine- or glutathione-conjugated, and therefore odorless, there exists a large potential pool of aroma that can be freed from β-lyase activity, such as by Irc7p (Roncoroni et al. 2011; Roland et al. 2016). Here, we observed a variable distribution of inactivating mutations in *IRC7* among the screened brewing strains and hybrids, as well as *IRC7* copy number variations between parents, hybrids and spore clones. These mutations have been demonstrated to directly influence wine thiol levels (Cordente et al. 2019). However, we only observed a minor, but positive, effect on beer thiol levels. A similar observation regarding lack of correlation between *IRC7* mutations and beer thiols levels was found in a recent study (Michel et al. 2019). It is possible that beer environment is not optimal for β-lyase activity, but that requires further clarification. Indeed, a recent study showed variation in amount of thiols released from supplemented glutathionylated and cysteinylated forms based on wort extract levels and fermentation temperature, but maximum release ratio for the bound forms remained below 0.5% and 0.1%, respectively (Chenot et al. 2021). Nevertheless, we succeeded in our goal of enhancing thiol release through breeding. Beer yeasts with enhanced β-lyase activity could help brewers heighten the flavour of popular beer styles such as “hazy” IPA, and/or reduce the cost impact of modern IPA hopping rates.

The use of genetically modified yeast for beverage production is still prohibited in most parts of the world, and the hybrid strains generated are considered genetically modified (Alperstein et al. 2020). However, the strains can be considered cisgenic or self-cloned, as no exogenous DNA is present in the cells and the Cas9 enzyme has only been used to create a DSB in the mating type locus (similarly to HO endonuclease). Hence, the strains are currently suitable for certain markets, including North America and Japan (Fischer et al. 2013). Regarding industrial suitability of the hybrids, previous yeast breeding studies have revealed that hybrid genomes may be unstable, and they can undergo substantial structural changes when repeatedly grown in a wort environment (Pérez-Través et al. 2014; Mertens et al. 2015; Krogerus et al. 2018). It is therefore vital that the long-term stability of the hybrids generated here is studied, through testing performance in beer fermentation and reuse over multiple yeast pitch generations. This is particularly important as polyploid strains have been shown to undergo chromosome losses during stress adaptation (Selmecki et al. 2015; Scott et al. 2017; Krogerus et al. 2018). Furthermore, suitability of these strains in combination with different hop varieties for production of aroma-forward beers has not yet been explored and may yield further insight into the aroma-enhancing potential of these yeasts.

In conclusion, our study confirms that CRISPR/Cas9-aided mating-type switching can be applied to homothallic aneuploid industrial yeast strains, and the switched strains can be readily mated to form hybrids. This allows for the rapid breeding of brewing strains, and overcomes the bottleneck caused by their sterility and polyploidy. The brewing hybrids constructed here exhibited heterosis across a variety of traits, including fermentation performance and aroma formation. Our results corroborate previous research highlighting the power of yeast breeding for strain development.

## Acknowledgements

We thank Aila Siltala, Niklas Fred, Eero Mattila and Ronja Eerikäinen for technical assistance, Dominik Mojzita for preparing the Cas9 plasmids, and George van der Merwe for sharing genome sequencing data.

## Availability of data and material

The Illumina reads generated in this study have been submitted to NCBI-SRA under BioProject number PRJNA740182 in the NCBI BioProject database (https://www.ncbi.nlm.nih.gov/bioproject/).

## Authors’ contributions

KK: Conceived the study, designed experiments, performed experiments, analysed all data, wrote the manuscript.

EF: Designed experiments, performed 400mL wort fermentations, edited the manuscript.

NR: Performed thiol analysis, edited the manuscript.

BG: Conceived the study, designed experiments, edited the manuscript.

RP: Conceived the study, designed experiments, edited the manuscript.

All authors read and approved the final manuscript.

