## Supplementary Figures for "Efficient breeding of industrial brewing yeast strains using CRISPR/Cas9-aided mating-type switching"

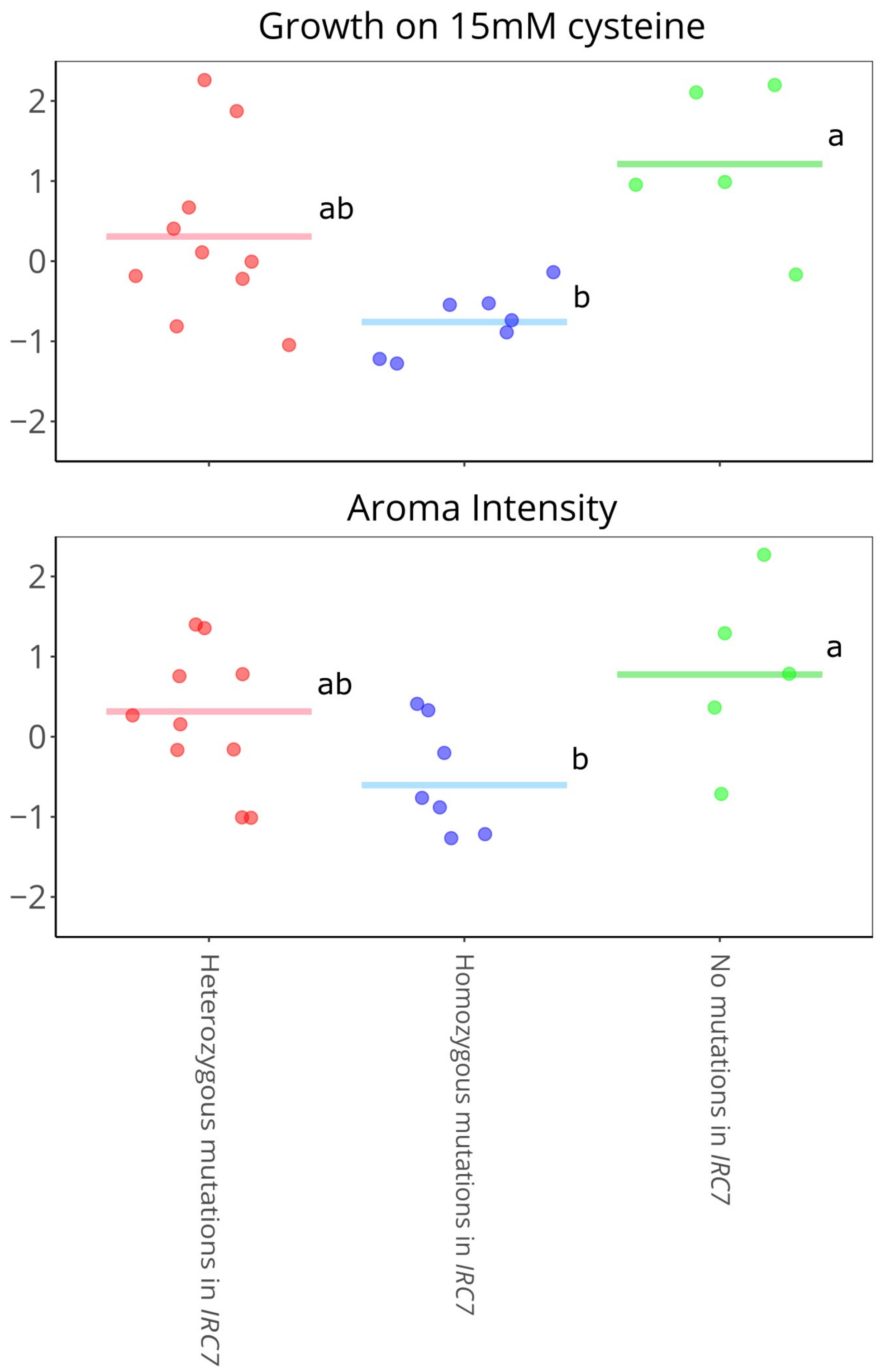


**Supplementary Figure S1** - Comparison of *S. cerevisiae* strains homozygous for the *IRC7* long allele. Growth on 15mM cysteine and aroma intensity in wort fermentations supplemented with Cys-4MMP was compared between strains harbouring either heterozygous, homozygous or no inactivating SNPs in *IRC7.* Groups with different letters differ significantly ($p < 0.05$).

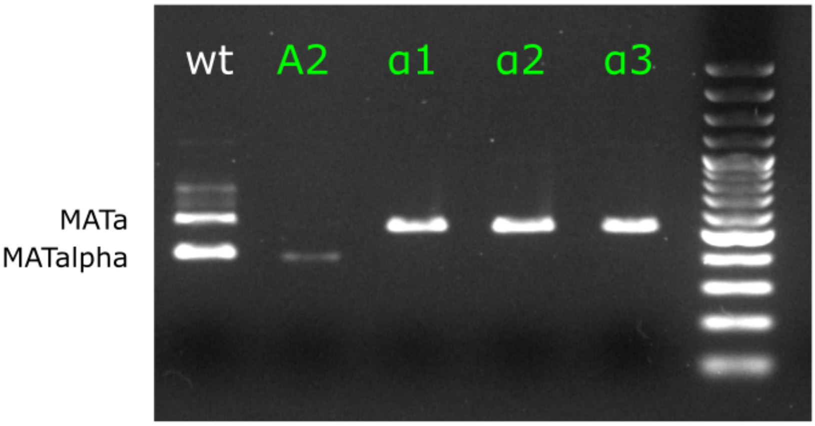


**Supplementary Figure S2** - Successful mating-type change as determined by PCR. The wild-type (wt) strain is heterozygous for the mating type locus. Transformant A2 has been transformed with a Cas9 plasmid targeting *MAT***a**, and produces only a band for *MAT*α. Transformants α1-3 have been transformed with a Cas9 plasmid targeting *MAT*α, and produces only a band for *MAT***a**. Primers used: MAT-R, MATa-F and MATα-F.

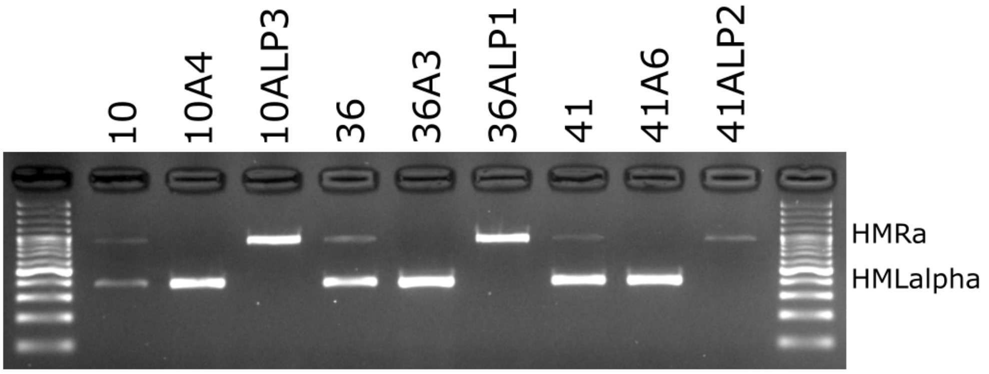


**Supplementary Figure S3** - *HMR***a** and *HML*α are simultaneously deleted in the transformed strains. Wild-type strains 10, 36 and 41 produce bands for both *HMR***a** and *HML*α. A transformants (Cas9 plasmid targeting *MAT***a**) produce bands only for *HML*α. Alpha transformants (Cas9 plasmid targeting *MAT*α) produce bands only for *HMR***a**.

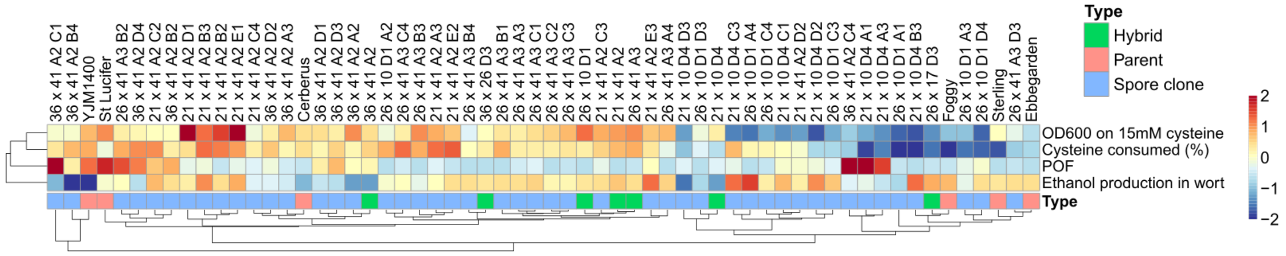


**Supplementary Figure S4 -** Phenotypic variation among the parent, hybrid, and spore clone strains. The heatmap is colored based on Z-scores (blue to red).

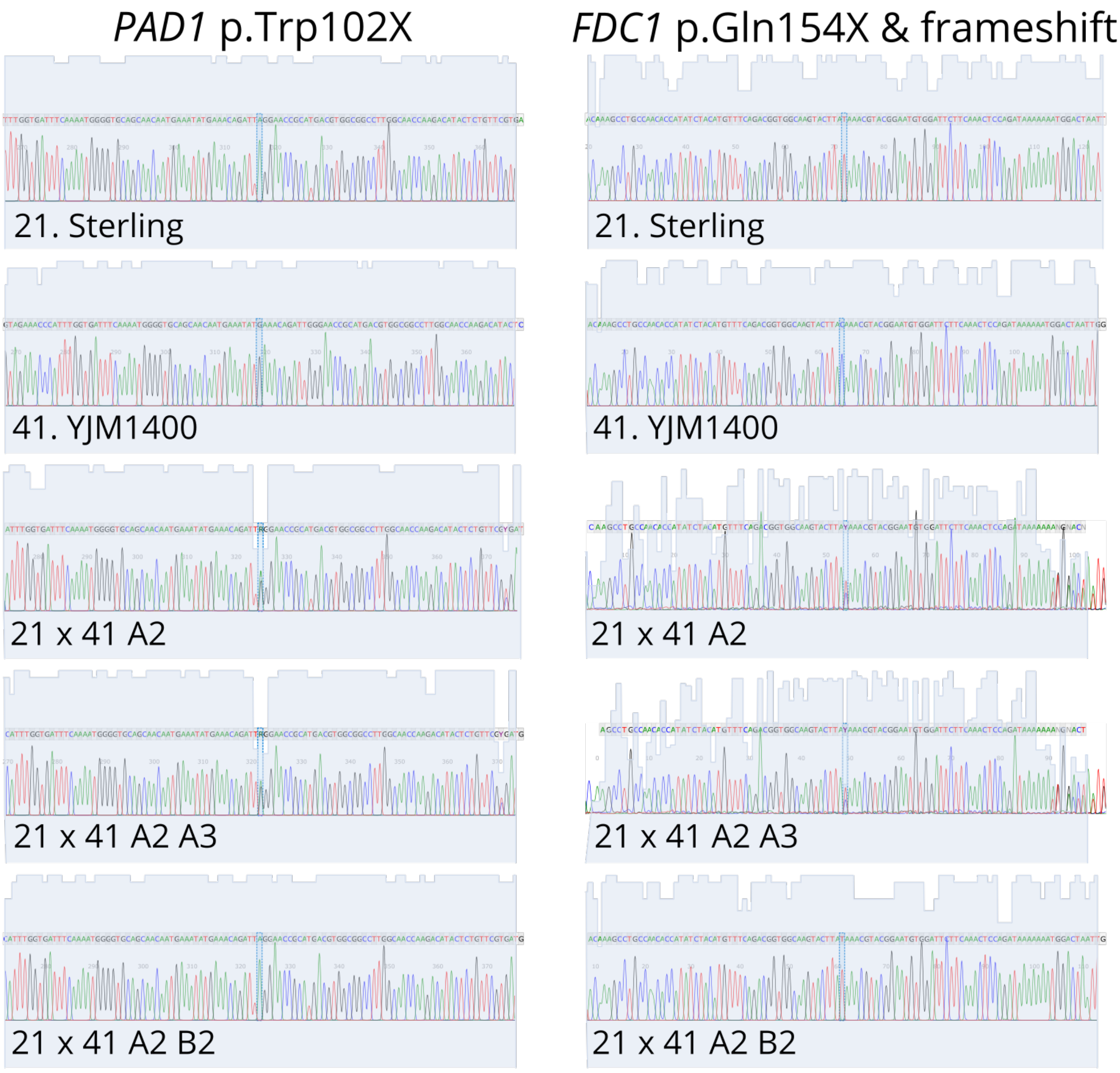


**Supplementary Figure S5** - Inactivating mutations in *PAD1* and *FDC1* among parent, hybrid, and selected spore clone strains. Strains 21 × 41 A2 and 21 × 41 A2 A3 are heterozygous for the inactivating mutations, while 21 × 41 A2 B2 is homozygous.
